# The Egg-Counter: A novel microfluidic platform for characterization of *Caenorhabditis elegans* egg-laying

**DOI:** 10.1101/2023.09.01.555781

**Authors:** Stephen A. Banse, Cody M. Jarrett, Kristin J. Robinson, Benjamin W. Blue, Emily L. Shaw, Patrick C. Phillips

## Abstract

Reproduction is a fundamental process that shapes the demography of every living organism yet is often difficult to assess with high precision in animals that produce large numbers of offspring. Here, we present a novel microfluidic research platform for studying *Caenorhabditis elegans’* egg-laying. The platform provides higher throughput than traditional solid-media assays while providing a very high degree of temporal resolution. Additionally, the environmental control enabled by microfluidic animal husbandry allows for experimental perturbations difficult to achieve with solid-media assays. We demonstrate the platform’s utility by characterizing *C. elegans* egg-laying behavior at two commonly used temperatures, 15 and 20°C. As expected, we observed a delayed onset of egg-laying at 15°C degrees, consistent with published temperature effects on development rate. Additionally, as seen in solid media studies, egg laying output was higher under the canonical 20°C conditions. While we validated the Egg-Counter with a study of temperature effects in wild-type animals, the platform is highly adaptable to any nematode egg-laying research where throughput or environmental control needs to be maximized without sacrificing temporal resolution.

## Introduction

The timing and extent of reproduction is the fundamental property underlying the biodemography of all organisms and underlies most processes—from evolution to aging—that shape biological function and diversity^1^ . For oviparous species that do not invest in parental care, choosing when and where to lay eggs is the final opportunity for a mother to influence the evolutionary fitness of her progeny^2^. As such, the egg-laying circuitry is modulated in response to sensory inputs that inform egg-laying decision making. For example, in *Drosophilids*, where egg-laying behavior has been widely studied using both comparative and functional approaches, site suitability for egg laying is assessed using sensory inputs like olfaction and taste^3–7^. The decision-making process is not a reflexive response to sensory input though, and behavioral output is modulated by situational context^8–11^, circadian rhythms^12,13^, and longer time cycle inputs like seasons^14,15^. When the decision to lay an egg is made, the neural circuit that enervates the egg-laying muscles activates a coordinated series of muscle contractions to achieve oviposition.

In the model nematode *Caenorhabditis elegans*, the relatively simple hermaphrodite’s anatomy consisting of only 302 neurons^16^ and 959 total somatic cells^17^, coupled with a myriad of genetic^18^, electrophysiological^19^, genetically encoded calcium^20^, and optogenetic tools^21^, has facilitated the identification of the core nematode egg laying circuitry^22,23^. Egg laying in *C. elegans* is primarily controlled by the serotonergic hermaphrodite specific <u>n</u>eurons (HSNL and HSNR) and the cholinergic VC motor neurons (VC1-VC6) [For review see ^22^]. These two classes of neurons, along with the vulval muscles and a neuroendocrine cell (uv1)^24^, form a circuit that generates “bursty” egg-laying behavior that is modulated in response to a variety of environmental factors^25,26^. This simple behavioral circuit, with the associated experimental tools, provides a powerful system for studying circuit modulation in response to sensory inputs^26^.

While the fundamental processes in *C. elegans* egg-laying behavior are generally well characterized, it should be noted that much of the research to date has been limited by experimental constraints. Standard assays for directly measuring *C. elegans* egg-laying behavior use low-throughput microscopy-based recording of animals on solid media. These assays require keeping behaving animals within an imaging field of view using motorized stages (worm trackers)^25,27,28^, or by physically restricting animal movement in static setups^29^. The physical requirements imposed by these approaches typically limit throughput to one animal per imaging rig. The low throughput of these imaging systems, and their relatively high cost, make it difficult to scale experiments. Additionally, longitudinal imaging of animals on solid media makes experimental modulation of the environment difficult, with no easy means for changing feeding conditions, temperatures, or chemical stimuli. As such, much of the investigation of environmental effects on *C. elegans* egg laying has been constrained to extreme conditions in which behavior can be indirectly measured in binary terms (e.g., “on/fast” or “off/slow”). In these experiments the finer temporal patterns of egg-laying are ignored, and the number of eggs retained in the uterus, the developmental stage of laid embryos, or the incidence of internal egg hatching, are used to infer if egg laying was slower or faster than normal^34–39^. This has left more nuanced influences on behavior relatively unstudied.

Here we present a microfluidic Egg-Counter that enables simultaneous monitoring of egg-laying behavior of up to 32 *C. elegans* hermaphrodites for long duration experiments (e.g., 48 hours) while providing precise temperature control (±0.1–0.2°C) across the permissive *C. elegans* growth temperature range. Using the Egg-Counter, we measured egg-laying at 15 and 20°C, two standard temperatures for *C. elegans* research^40^. We observed that behavior in our microfluidic chip is similar to behavior in traditional solid-media studies, with eggs laid in bursts. Additionally, reproductive output appears higher at 20°C, the standard growth temperature, than at 15°C, consistent with *C. elegans* solid-media assays^41,42^. In total, our characterization of temperature effects on *C. elegans* egg-laying with the Egg-Counter demonstrates the microfluidic research platform’s utility for studying the modulation of a powerful model behavioral circuit.

## Results

### The Egg-Counter Chip

The microfluidic Egg-Counter chip was designed to simultaneously house up to 32 animals in individual growth arenas (Figure 1A). The growth arenas are arrayed in two groups of sixteen, with independent perfusion inlets and upstream distribution networks to enable side-by-side comparisons of genotypes or perfusion conditions. The distribution networks end at a loading chamber^30,31^ just upstream of each growth arena that has been sized to sequentially trap, then load, L4 staged animals (Figure 1B). After animals are loaded, they are housed for the duration of the experiment in growth arenas containing an array of pillars that promote sinusoidal “crawling” movement similar to animals on solid media^32,33^. This is important because *C elegans* egg laying is tightly coordinated with physical movement^28,43,44^, and free swimming animals are prone to slow, or stop, egg laying and die through internal hatching^45,46^. The arenas’ exits feature filters sized to allow eggs to pass in the buffer outflow while retaining adults. After exiting the arenas, eggs travel in separate outflow channels that are brought together in parallel to pass through an imaging zone (Figure 1C). The imaging zone enables eggs laid in all 32 arenas to be observed in one location, greatly increasing the potential throughput over traditional worm-tracker assays. Using a flow-through microfluidic device also enables delivery of a consistent diet and experimental stimuli, while removing animal waste and secretions^47,48^. Additionally, the physical arrangement allows for future addition of off-chip components for modulation of perfusate for temporal presentation of diets and stimuli as has been done for other *C. elegans* microfluidic devices^32,33,49–51^.

**Figure 1.**
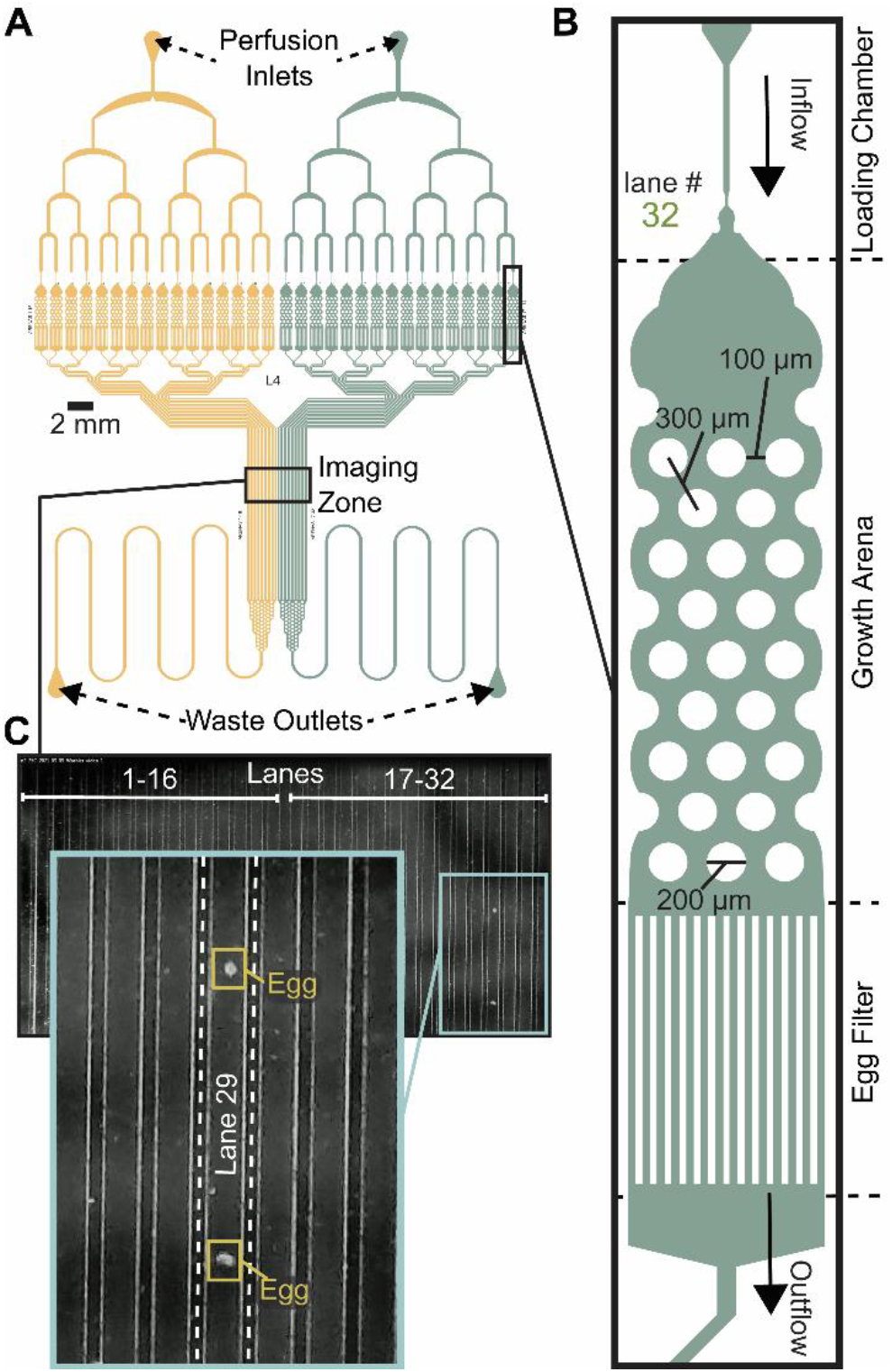
Egg-Counter chip design features. (A) Schematic of the Egg-Counter chip. The chip features two independent arrays (yellow and green) that begin with a perfusion inlet for buffer/food flow during an experiment. The fluid flows through a distribution network before entering 16 parallel animal Growth Arenas. The exit channels from each Growth Arena remain separate until after the Imaging Zone when the channels merge and continue to the Waste Outlet. Additional channel length prior to the Waste Outlet adds system resistance to minimize changes in flow rate during experiments. (B) Each of the Growth Arenas consist of three sequential features; a Loading Chamber that uses a hold and push method^30,31^ to facilitate loading animals, the actual arena featuring a hexagonal pattern of pillars that promotes plate-like sinusoidal movement^31–33^, and an Egg Filter that retains adults while allowing eggs to pass into the outflow. (C) The individual outflow channels are brought together in a parallel arrangement in the imaging zone. Shown is a representative image of the imaging zone at a time when two eggs laid in Growth Arena #29 were detected.

### The Egg-Counter platform

*C. elegans* egg laying is sensitive to the chemical environment, with variables like food concentration, CO_2_ level, and osmolarity impacting egg-laying behavior^29,54–56^. The chemical components of the experimental environment are therefore standardized prior to entry into the microfluidic device in the Fluid Control portion of the platform (Figure 2A). For example, the uniformity of the environmental buffer and bacterial food suspension is maintained throughout the 48-hour experiments using a stir-bar and stir-plate and supplementation with bacteriostatic antibiotics.

**Figure 2.**
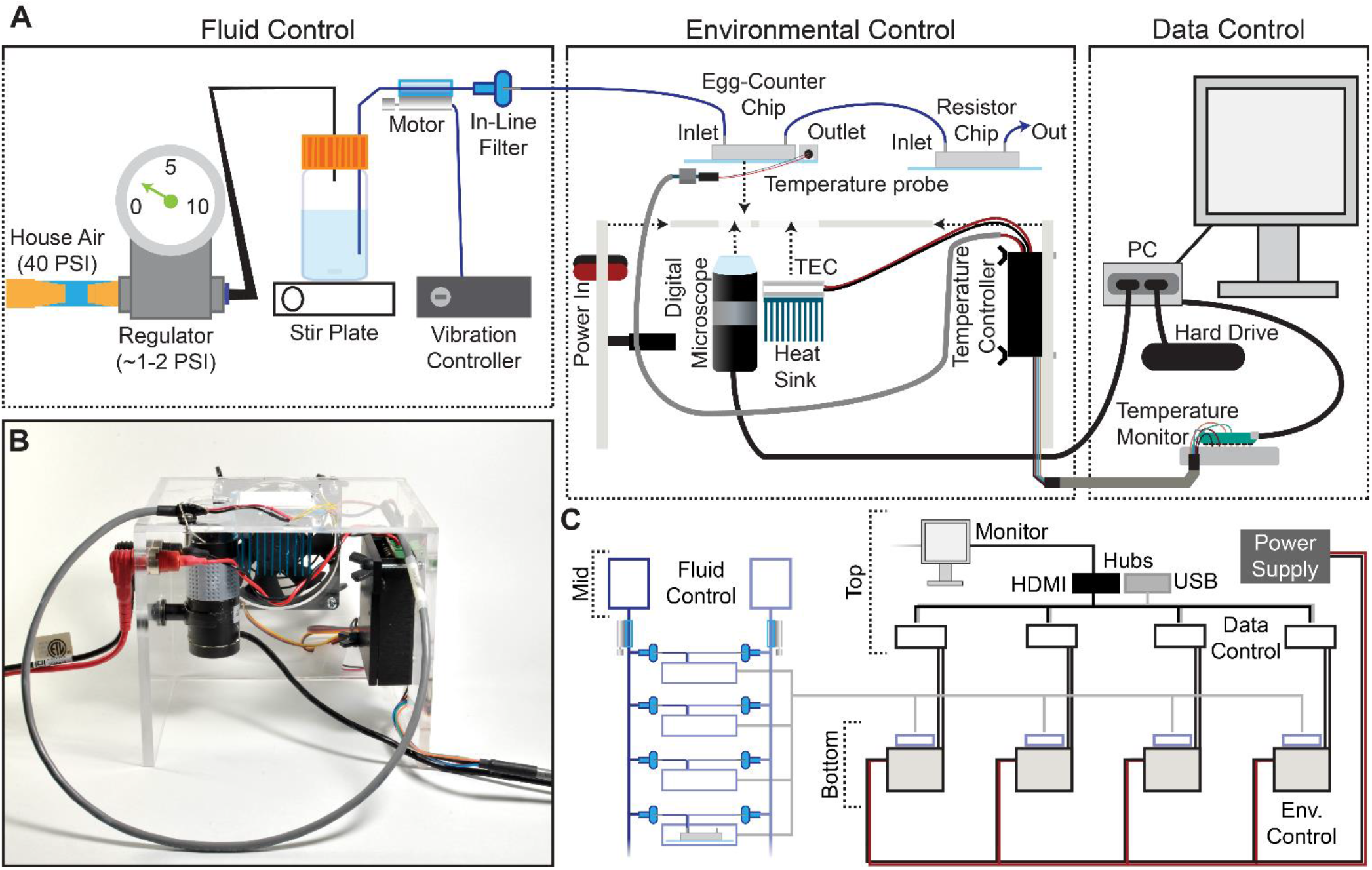
Egg-Counter platform. (A) The Egg-Counter platform comprises three control systems: Fluid Control, Environmental Control, and Data Control. Fluid flow is driven by pressurizing the air in the buffer bottle to ∼0.5 PSI. The buffer containing suspended bacteria is then driven through tubing with an attached vibration motor that maintains the bacterial suspension. Prior to entering the Egg-Counter chip, the fluid passes through a 5 µm filter. Egg-Counter chips are mounted on the Egg-Counter platform for environmental control. A custom acrylic platform holds a digital microscope, thermoelectric cooler (TEC), heat sink, fan, and temperature controller. The chip is mounted directly on the TEC with thermal paste to maximize thermal contact. A temperature probe mounted on the glass slide under a PDMS cover, with potential air pockets displaced by thermal paste, feeds temperature data to the temperature controller. The Temperature Controller reads the temperature from the glass surface that the animals contact and controls the TEC to maintain the setpoint temperature. Data control is performed by a fanless computer and a Temperature Monitor. The computer takes experiment parameters from the user, communicates with the Temperature Monitor (which sets the temperature on the temperature controller), and starts the experiment. The PC monitors a live feed from the digital microscope and saves all frames with detectible changes. The microcontroller maintains communication with the temperature controller and receives temperature data at 5-second intervals, ultimately transferring the temperature logs to the PC. (B) An assembled Egg-Counter platform (minus microfluidic chip and fluid tubing). (C) The Egg-Counter was designed to be scalable while minimizing the footprint. Two shelves above a counter arrangement, with the top shelf housing a single monitor (for up to 8 devices), the Data Control electronic components, and custom power supplies each capable of powering 4 devices. The middle shelf houses the Fluid Control elements. Each bottle can feed 6-8 inlets (3-4 chips) and bundling the tubing between the chips and the bottle enables a single vibration controller to service multiple lines. The bottom shelf or countertop houses the environmental control platform, which is built to maintain the electronics in an elevated position. This configuration helps to separate the pressurized fluid and the electronic components.

During preliminary experiments we learned that the relatively slow linear velocity in the tubing connecting the food/buffer bottle to the microfluidic chip facilitated bacterial sedimentation prior to entry into the microfluidic device. We therefore added a neoteric line-agitation device that uses vibration motors and a custom controller to keep bacteria in suspension before entering the Egg-Counter chip (Figure 2A; See online collection^52^). To avoid confounding effects from vibrations^57^, the motor was attached between the buffer bottle and the inline filter (Figure 2A), and the tubing was physically tethered just upstream of the Egg-Counter chip to dampen vibrations before reaching the animals.

Because environmental variables other than chemical composition (e.g., vibration^57^, light^58^, and temperature^59^) have profound effects on *C. elegans*, we designed the Environmental Control system (Figure 2A, B) to minimize confounding variables. For example, the platform’s constant video monitoring of the imaging zone (Figure 1A and C) uses a digital microscope and light source located underneath the microfluidic chip. The growth arenas sit on an opaque surface offset from the imaging zone to avoid direct illumination. The platform regulates temperature using an embedded thermoelectric cooler (TEC) that sits directly under the growth arenas while air pockets are displaced with thermal paste. During experimental runs, a temperature probe mounted on the Egg-Counter chip’s glass surface in a mock PDMS chip provides real-time temperature data to the Temperature Controller (Figure 2A).

The materials needed to construct a full Egg-Counter rig are relatively inexpensive, with a total parts cost under $1,400. The actual cost would depend on the number of devices built because the system is designed to be scalable (Figure 2C). As such, the actual costs range from ∼$1,200 to 1,375 each for a complete Egg-Counter and computer (see parts list in the online collection^52^). In comparison, it would cost more than $15,000^60^ in parts to build an open-source worm tracker for traditional egg-laying assays before accounting for the necessary computer and environmental control. We conclude that based on the cost per animal studied, the Egg-Counter can provide a 400-fold or greater increase in throughput over traditional approaches.

### Egg-Counter temperature control

To avoid the need for expensive incubators, the Egg-Counter controls the experimental temperature at the point of animal husbandry. We integrated the temperature control and monitoring into the experimental control software that is run on a mini-format PC that directly connects to the digital microscope and a custom-built Temperature Monitor (Figure 3A, B). The temperature monitoring unit acts as a mid-level coordinator, providing the voltage setpoint for the desired temperature to the temperature control unit and logging the experimental temperature at five second intervals for the duration of the experiment. We present representative temperature profiles from two independent runs on the same rig for each of the five temperature settings used (Figure 3C). We observed that the recorded temperatures for a rig are consistent during and between experimental runs, with clear separation between temperature setpoints. Comparisons between temperatures suggest that there is a relationship between temperature setpoint and the spread in observed temperatures, with the standard deviation in temperature increasing from 0.098°C at 15°C to 0.20°C at 25°C (*p*<.0001). Full temperature recordings, and summaries of the between-rig offsets are available in an online collection^61^, along with the temperature control software.

**Figure 3.**
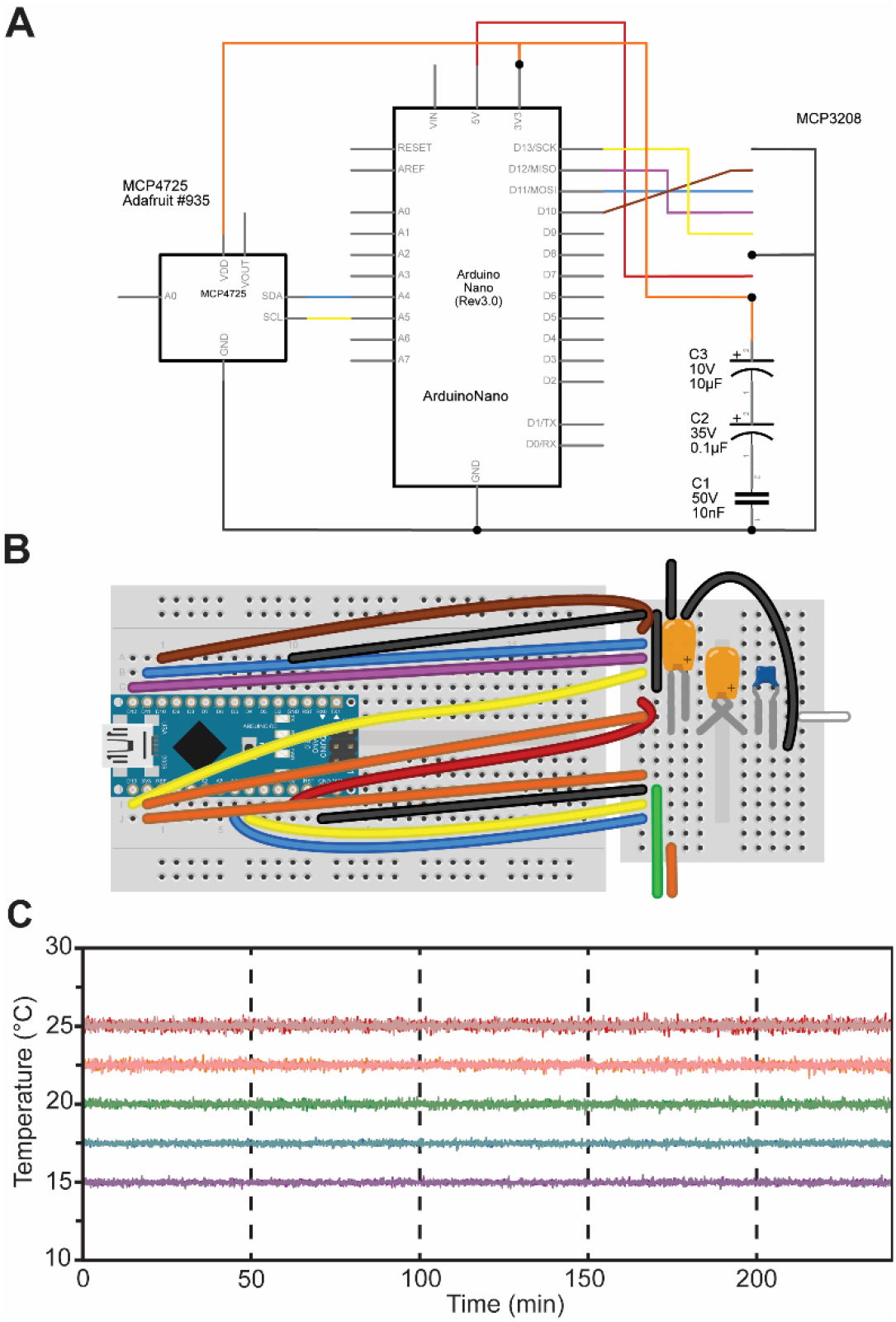
Egg-Counter Temperature Monitor. (A) Electrical schematic and (B) breadboard diagram for the Egg-Counter Temperature Monitor. Parts list and necessary software are available in online supplements ^52^ and ^53^ respectively. (C) Representative temperature data from two experimental runs (shown in light and dark shades) at five temperature conditions covering the standard growth range of *C. elegans* (15°C purple, 17.5°C blue, 20°C green, 22.5°C orange, and 25°C red).

### Egg-Counter experimental control

To run the experiments, we created software for automated experiment control (see online collection ^53^). The software allows the user to select the temperature profile for the experiment, duration of the experiment, imaging settings and enter relevant metadata through a graphical user interface (GUI) (Supplemental figure 1). The software then initiates the experiment, sets the temperature, and launches the egg detection software that records the egg-laying data. The egg monitoring software (Supplemental figure 1; online collection^53^) connects to a digital microscope that is focused on the “imaging zone” (Figure 1) to monitor for eggs. It proved impractical to record 48-hour long movies at ∼10 fps, so the software continually monitors the video feed and compares sequential frames to detect movement (Supplemental figure 2). When moving objects are detected, the relevant image frames and associated metadata are logged. To facilitate downstream analysis, the potential egg containing frames are saved as a movie file and all associated frame meta data are saved in a spreadsheet at 12-hour intervals.

To process the saved videos to determine the timing of egg-laying events, the videos were put through an annotation pipeline to determine which frames contained eggs, which frame represented the first detection of each egg, and which objects represented clusters of eggs. To annotate eggs within the collected images, we initially attempted automated egg identification using random forest classifiers. While the automated approaches were intriguing, the limited feature complexity of oval or round “egg” objects did not provide enough detail to easily train an automated system to achieve accuracies better than 80 to 90% when cross validated by human assessment. We therefore created a computer aided human annotation pipeline in which multiple parameters from an automated contour analysis were used to identify potential egg objects for annotation (Supplemental figure 3; online collection^53^). Our software presents auto-detected egg-like objects in a GUI for the user to annotate (see online collection^53^). After annotation, the software generates a list of eggs laid in each lane along with the associated time of first detection. To validate this approach, we had different experimenters annotate the same sample datasets to determine the repeatability of the annotation process. When two experimenters annotate the same dataset there is a high concordance between the two sets (see online supplemental collection^61^), suggesting that our computer aided annotation provides reproducible results.

### Temperature affects C. elegans egg-laying

To quantify egg-laying behavior in our microfluidic Egg-Counter we raised animals in liquid culture at the standard growth temperature of 20°C. Animals were then transferred to the microfluidic device at the L4 stage to begin 48-hour experiments. At the end of the experiment the movies were analyzed and annotated as described above.

When we plotted the temporal density of laid eggs from individual animals, we observed a variety of potential changes in egg-laying patterns between the tested temperatures (Figure 4A). For example, temperature appeared to change the time at which the first egg laid was detected (Figure 4A). When we quantified the time to egg-laying onset (Figure 4B) we found that temperature had a significant impact (*p*<.0001), with the mean onset time accelerating from ∼13.2 hours at 15°C to ∼9.9 hours at 20°C (*p*<.0001). This result is roughly consistent with the published difference in developmental rate observed for these experimental temperatures, although published developmental-stage-specific deviations in temperature effects may suggest that a strong correlation between observations unexpected ^42,62,63^. Regardless of that caveat, the observed difference in onset time demonstrates that the animals were responsive to the experimental temperature regimes. Quantifying the number of eggs laid during the 48-hour experimental period showed that temperature affects reproductive output. Similar to plate-based assays^41,42^, egg-output is higher at the standard lab temperature of 20°C (100.6 eggs) than at 15°C (51.7 eggs at *p*<.0001) (Figure 4C). The total number of eggs laid within the experimental period is determined by both the time of onset and the egg-laying rate (ELR) after onset. We therefore determine the ELR, defined as the slope of a linear fit to the post onset egg-laying trajectory, for each animal (Figure 4D). We observed that the ELR after onset changed with temperature (*p*<.0001), with the rate effectively doubling from 1.5 eggs/hour at 15°C to 2.9 eggs/hour at 20°C. (*p*<.0001). The fact that temperature changes the observed ELR suggests that the observed differences in reproductive output were not due solely to effects on the initiation of egg laying, and that temperature continues to affect egg laying in adulthood.

**Figure 4.**
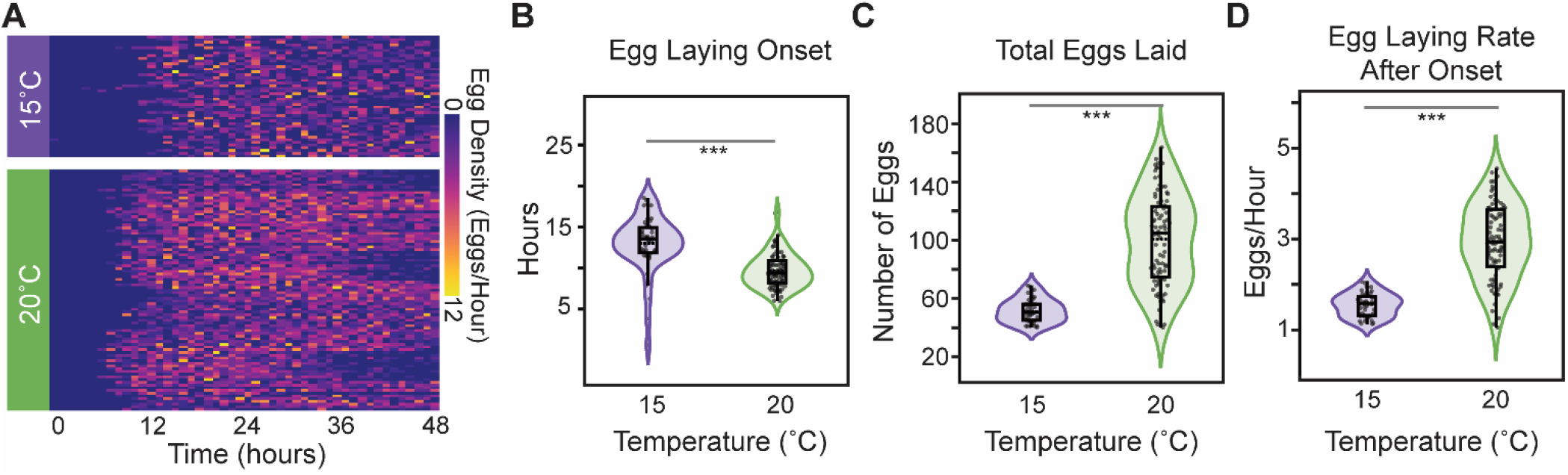
Temperature effects on egg laying. (A) Heat map of the number of eggs laid in 1-hour windows over the course of the 48-hour experiment at the two tested temperatures, (B) The egg-laying onset time at the five experimental temperatures. (C) The total number of observed eggs laid during the 48-hour experiment. (D) The egg-laying rate, defined as the slope from a linear regression for cumulative number of eggs laid versus time. All panels use the same color key to distinguish the two temperatures (15°C purple and 20°C green). Presented statistical analyses (*** *p*<0.001) are nonparametric Wilcoxon comparisons.

### C. elegans exhibits bursty egg-laying in the Egg-Counter

We next sought to determine if egg laying in the Egg-Counter was similar to the “bursty” behavior on solid media, with periods of inactivity punctuated by relatively rapid laying of eggs^25,26,64^. The bursty nature has been postulated to arise from the transition between neuromuscular states of active and inactive egg-laying. This results in a bimodal distribution of inter-egg intervals. Within this interpretation, the egg-laying circuit is in the active state for a matter of minutes, and during that period there can be a burst of egg-laying activity marked by inter-egg intervals that are seconds to minutes long (within-bout-intervals [WBI]; Figure 5A). In contrast, residency time in the inactive state results in second class of longer duration inter-egg intervals, as those intervals represent the time between the last egg of a bout and the first egg of the following bout (inter-bout-intervals [IBI]; Figure 5A). To determine if our data exhibited similar patterning, we plotted the cumulative egg counts at each time point from the canonical “control” temperature of 20°C to generate the egg-laying trajectories for animals under control conditions (Figure 5B). We see the expected pattern of egg laying in our microfluidic device, with the egg-laying trajectory exhibiting periods of inactivity followed by bursts of egg laying (Figure 5B inset), which provides qualitative support to the hypothesis that our data is patterned similarly to the behavior on solid media.

**Figure 5.**
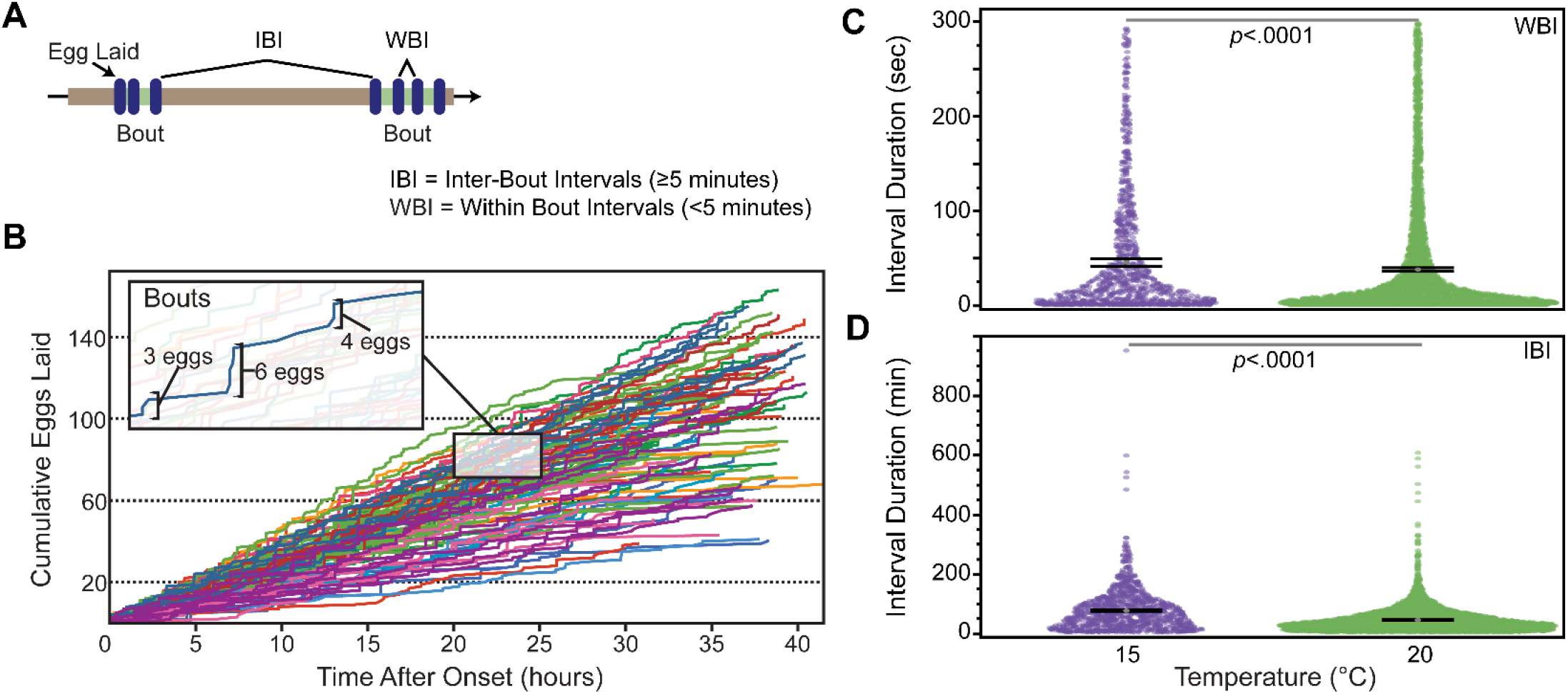
Egg-laying trajectories in the Egg-Counter. (A) The distribution of egg laying events in time is distributed such that two inter-egg intervals exist. (B) Egg-laying trajectories from the time of egg-laying onset for animals at 20°C degrees. Shown are data from 12 different Egg-Counter chips, with the traces colored by experimental chip. The figure inset shows a 5-hour segment with one trajectory highlighted to illustrate the observed “bursty” pattern of periods with no eggs laid, punctuated by bouts of active egg laying. In the example, bursts of 3, 6, and 4 laid eggs are noted. (C) The distribution of within-burst (WBI) intervals (*N* = 1,103 at 15°C and 4,702 at 20°C), and (D) the distribution of inter-burst (IBI) intervals (*N* = 1,126 at 15°C and 4163 at 20°C) with their respective mean and 95% confidence interval is shown.

To further evaluate the pattern of egg-laying in the Egg-Counter, we analyzed the pooled intervals from the animals at the two tested temperatures. To separate inter-burst intervals (IBI) from within-burst intervals (WBI), we set out to determine an appropriate cut-off to separate the two classes of intervals. Previous published analysis of egg-laying behavior on solid media had determined an average within bout interval of ∼20s and an average inter-bout interval of ∼20 minutes^25^. Additionally, it has been shown that plotting the log-tail interval distributions provides a visualizable transition between the two classes of intervals^25,64^. We therefore generated the log-tail interval plots for our data at 15°C and 20°C (Supplemental fig 4). As has been seen in previous analyses, there was a clear change in slope between two classes of interval. Using this analysis we settled on a cut-off of 300 seconds, which is generally consistent with published data at 20°C^25,64^. We next separated our pooled intervals using the selected cut-off and observed a statistically significant (*p*<.0001), difference in the mean WBI at 20°C (38.2 seconds) and 15°C (45.6 seconds) (Figure 5C). In contrast to the ∼19% difference in mean WBI, we observed a larger difference in the IBIs (*p*<.0001), with the mean IBI at 20°C (46.2 minutes) increasing ∼68% at 15°C (77.6 minutes). We next evaluated the distribution of intervals between WBIs and IBIs at the two temperatures. We observed a negligible temperature dependent difference in the distribution between the two classes of intervals, with the fraction of intervals representing inter-bout intervals increasing from 47% at 20°C to 50% at 15°C (*p*= .0035).

## Discussion

The Egg-Counter is a novel high precision platform for studying egg laying in *Caenorhabditis* nematodes that is built around a novel microfluidic chip (Figure 1) housing up to 32 individual animals in separate Growth Arenas under precise environmental control. We find that individuals raised in this environment exhibit “normal” movement like that observed on solid media^32,33^. The Growth Arenas are divided into two separate arrays to enable side-by-side comparison of genetic backgrounds, food levels, or environmental stimuli. The flow-through chip enables experimental control of feeding and environmental standardization, while flushing embryos out as they are laid. The outflow from each arena remains separate until after all out-flow channels are brought into a parallel configuration and pass through a central imaging zone for simultaneous monitoring for eggs from all arenas in the chip. The microfluidic chip is mounted on a custom temperature control and imaging device that connects with a control computer for image acquisition and a custom temperature logger (Figure 3). The system is scalable in groups of four (Figure 2), with each grouping sharing a common power supply being capable of housing up to 128 animals (see online collection ^52^). In the online supplemental collection, we provide a detailed parts list, experimental procedures, and open-source software for the platform^52,53,61^.

### The benefits of the microfluidic Egg-Counter

Traditional egg-laying assays for *Caenorhabditis* nematodes are conducted on solid media using video-capture to determine the timing of egg-laying events. The need to confine individuals within the microscopic field of view, or use motorized stage worm trackers, greatly limits the throughput of traditional assays. Additionally, the use of solid media makes experimental modulation of the environment difficult. These limitations have confined most research to shorter assay periods, required temperature-controlled incubators or rooms, and limited the number of animals studied. In contrast to traditional assays, experiments using our Egg-Counter platform allow simultaneous recording of egg laying from multiple animals, enables controlled experimental temperature on a benchtop, and allow for multi-day experiments. The fine temporal resolution enables analyses of temporally patterned data that is currently only available through low-throughput systems. Additionally, while our proof-of-principle experiments did not vary the environment within experimental runs, the flow through system and temperature control rig can provide reproducible environmental perturbations like cycling food or temperature regimes, opening avenues of research not previously available (see ^65–67^ for examples of in-chip environmental perturbations). As such, the Egg-Counter provides several significant benefits over traditional solid-media assays.

### Microfluidic alternatives to the Egg-Counter

While the Egg-Counter has many benefits over traditional egg-laying assays, other microfluidic devices can be used to measure reproductive output of *C. elegans*. For example, a microfluidic device designed to follow reproductive aging has been used to follow the progeny production from multiple animals simultaneously through an imaging system similar to ours^68^. Despite the similarity, the previously published device was incapable of following egg laying, and instead used a filtration setup to separate the thinner L1 larvae from adults. The requirement for embryos to complete embryonic development and hatch as L1s before exiting the housing arena adds a delay of 8-18 hours from the time of egg laying^41^, rendering the assay blind to the egg laying behavior itself. Alternatively, a second microfluidic device has been developed to follow egg laying by catching laid eggs in a “basket” for quantification^65,69^. Unlike our device and the reproductive aging chip^68^, the animals are confined for the duration of the experiment, and the important regulatory connection between animal motility and egg laying is disrupted^65,69^. A third microfluidic device has been developed to follow egg laying induced by electrical fields^67,70^. While electrically induced egg-laying represents a novel assay, and has many potential uses in shorter duration experiments, the device also uses a restrictive small animal holding chamber that would greatly restrict normal sinusoidal movement. Additionally, the typical electrically stimulated egg laying experiment is much shorter (10 minutes) than for the Egg-Counter and the other two published relevant microfluidic devices. In total, the Egg-Counter presents a novel set of capabilities and strengths for assaying nematode egg laying in a microfluidic environment.

Of course, every different approach will have its own strengths and weaknesses. The Egg-Counter is especially well-suited for assessing patterns of reproduction at high temporal resolution, which for instance, allows for inter-individual differences to be measured with high precision. In practice, we find that the limiting factor in overall throughput is ensuring that single individuals are loaded into each chamber. Double loading or empty wells reduce the yield of individual measures and are an inherent limitation of the current loading design. Balancing size constraints of filtering larval worms from intact eggs remains a broader design challenge for the *C. elegans* microfluidics field.

### Temperature effects on egg-laying

To validate our platform, we collected data at two commonly used *C. elegans* rearing temperatures. We observed that the patterning of egg-laying was “bursty”, with long-periods of inactivity punctuated by groups of egg-laying events in quick succession, as is seen in traditional solid-media studies. Analysis of the intervals gives a similar bimodal distribution, with a mean in-bout interval of 38.2 seconds and a mean between-bout interval of 46.2 minutes. This compares to ∼20s^25^ and ∼20 minutes respectively on solid media at 20°C^25^. This suggests that while the egg-laying patterning is broadly distributed in the same bimodal manner in our device and in traditional assays, the overall rate of egg-laying is lower in the microfluidic device. We suspect that this is due to the relative concentrations of food in the two situations. Previous work has shown that the rate of egg-laying changes in response to food concentration^71^. We conclude that at a gross phenotypic level, animals in the Egg-Counter exhibit “bursty” behavior like that on solid media.

### Summary

Here we present a novel microfluidic research platform for studying *Caenorhabditis* egg laying. We provide a microfluidic device design, a temperature control system, image acquisition software and processing software. Using this system, we demonstrate that temperature has a negligible effect on the inter-egg intervals during bouts of egg-laying. This contrasts with a significant change in the inactive periods between egg-laying bouts. Additionally, we observe a change in the distribution of intervals between within-bout and between-bout classes, suggesting that temperature has a small but significant effect on the number of eggs laid within a bout. The platform therefore was successful in characterizing behavioral patterning for a research model with a known neural circuit that can be modulated by sensory input. As such it should have applicability to many different studies.

## Experimental Procedures

We provide detailed materials and methods in an online collection^52,53^ that includes: (1) all necessary CAD files for fabrication of the microfluidic Egg-Counter and resistance chips, 3D printing the vibration controller box, and laser cutting the components for the platform enclosure, (2) a detailed parts lists with ordering information, (3) open-source software for acquisition, processing, and annotation of egg images, (4) experimental SOPs, and (5) the experimental data underlying the figures presented in this manuscript. While the online supplement provides detailed information, the experimental procedures in brief are as follows:

### Microfluidic design

For this project we designed two new microfluidic chips. The first, the Egg-Counter (Figure 1A, Supplemental figure 5 and online collection ^52^), is designed to house 32 animals in two arrays of 16 growth chambers. Each growth chamber features an array of 200 µm pillars with a midpoint-to-midpoint spacing of 300 µm as previously published^31^. The second, the resistance chip (Figure 2a, Supplemental figure 6 and online collection ^52^), is used in series with the Egg-Counter chip to increase the total resistance for the system. The increase in total resistance better aligns the operating pressure with the operating range of the pressure regulator, enabling finer control of perfusion speed, while minimizing negative effects due to variability in pressure regulation over the duration of an experiment as previously published^72^. Both devices were designed using Vectorworks 2013 Fundamentals (Nemetschek SE, Munich, DE). Photomask transparencies were printed at 25,400 dpi (“20k resolution”, CAD/Art Services Inc, Bandon OR, United States). Printable CAD files are available in the online collection ^52^.

### Microfluidic device fabrication

We fabricated microfluidic devices using standard soft lithography techniques^73,74^. In brief, molds were made using negative UV photoresist (SU-8 2025, Kayaku Advanced Materials, [formerly Microchem]) spun coat onto silicon wafers (10 s at 500 rpm, with a ramp of 100 rpm/s followed by 30 s at 2000 rpm, with a ramp of 300 rpm/s). The expected feature height was ∼40 µm. The Egg-Counter and resistor designs were fabricated with the same spin speeds and SU-8. After fabrication, the SU-8 feature heights were measured using a Dektak 6M stylus profilometer. The mean height of the molds used in this study was 38 µm, with a minimum measured height of 37 µm and a maximum measured height of 41 µm. Microfluidic devices were then fabricated using Polydimethylsiloxane (PDMS) (Sylgard 184, Dow Corning Corporation) at the recommended ratio of 10-elastomer:1-curing agent by weight. 1.25 mm holes were punched using biopsy punches to gain access to the patterned channels, and the PDMS devices were bonded to 50x75 mm glass slides by air plasma exposure (PDC-32G Plasma Cleaner, Harrick Plasma Inc.).

### Bacterial strains and growth

Nematodes in the Egg-Counter were fed a derivative of *Escherichia coli* strain NA22 (available from the CGC -https://cgc.umn.edu/strain/NA22). To facilitate selection during culturing, carbenicillin/ampicillin resistance was introduced into NA22 competent cells using pUC19^75^ (Addgene, product #50005) to generate strain PXKR1. The plasmid harboring *E. coli* were propagated and maintained through either ampicillin or carbenicillin (carb) selection. A single PXKR1 colony was selected to inoculate 1 L of LB broth supplemented with ampicillin (100 mg/liter). The cultures were incubated at 37°C for 16-18 hours in a rotating incubator (at 180 rpm). After decanting into macro-centrifuge bottles, the cultures were pelleted at 8000 rpm for 10 minutes. The supernatant was disposed of, and the pellets were vortexed in ∼50 mL of 1xM9 and combined. To remove cellular aggregates or debris, the *E. coli* M9 suspension was filtered using a 5 µM syringe filter (PALL). After filtration, spectrophotometer measurements of OD600 were taken across a dilution series of the concentrated food stocks to determine the cell density. The stock solution was then used to generate solutions of 2x10^9^ PXKR1 cells/ml in 0.5x M9 supplemented with Kanamycin (50 mg/L) and carbenicillin (50 mg/L) to prevent culture growth and bacterial contamination. Additionally, Tween-20 was also added (1x10^-4^ v/v) to prevent embryos from sticking within microfluidic channels. A stir bar ++and stir plate were used to keep the cells in suspension. For experiments at >20°C the 1-liter bottle was placed onto a heating stir plate to pre-warm the culture to the TEC temperature. Initial experiments at temperatures above ambient experienced frequent failure due to air bubbles that formed in the microfluidic chip. We hypothesized that the lower oxygen carrying capacity of the buffer upon warming in the device was causing oxygen to be driven out of buffer. Pre-warming the solution to match the TEC programmed chip temperature circumvented this issue, and all experiments above ambient included a heated stir-plate to normalize the temperature and dissolved oxygen prior to on-chip temperature regulation. Each 1-liter bottle had six access ports pulling from the bacterial solution and was therefore capable of supplying food to 96 arenas/individuals equal to 3 chips.

### Animal husbandry and worm preparation

All experiments were performed with the N2-PD1073 wildtype reference strain^76,77^. To prevent bagging and foster familiarity, worms were reared in a liquid culture. Standard hypochlorite protocols were applied to arrest L1 staged worms overnight at 20°C in NGM buffer (cholesterol omitted)^78,79^ containing 1mg/ml carb, 1 mg/ml strep, and 1 mg/ml of nystatin. The next day the L1 density was calculated, and 500 worms were transferred to a fresh solution containing food. The synchronous population of L1s were given 5x10^9^ cells/ml of PXKR1 in NGM supplemented with 1 mg/ml carb, 1mg/ml strep, 1 mg/ml nystatin, and 0.5mg/ml cholesterol. The worms were then aged to the L4 stage in a 20°C incubator (48hrs). All of the above steps occurred in a 15 ml conical tube constantly inverted by rotator. Only PXKR1 liquid culture pre-conditioned worms were used in the Egg-Counter experiments. On the day of the experiment approximately 32 worms were pipetted into the entrance port of the Egg-Counter chip. Prior to loading the chip was degassed in a vacuum desiccator for 5 minutes to remove air bubbles from the PDMS. It was then treated with a surfactant 5% Pluronic F127 in dH_2_O for 40 min and rinsed with 1X M9. A syringe containing the experimental perfusate (2x10^9^ cells/ml of PXKR1 in 0.5xM9, kanamycin 50 mg/L, carbenicillin 50mg/L, 1x10^-^4 v/v Tween-20) was inserted and hand pressure was used to push the worms into individual arenas. Loading efficiency varied but ideally each activity arena contained a single worm.

### A note on larval growth

Interestingly, we found that the experimental design required growth in liquid culture at food concentrations similar to those experienced during microfluidic experimentation. With the caveat of starting at a higher concentration to account for adequate feeding during the 48-hour incubation period, hence 5x10^9^ rearing concentration. When the larval rearing conditions did not match the microfluidic experimental conditions, nearly 100% of the animals died by internal hatching of embryos^80^. Developmental conditions affecting adult behavior are not uncommon, and previous work has shown that adult physiological outcomes can be shaped by larval experience. For example, temperature is typically negatively associated with nematode lifespan, with *C. elegans* living longer at cooler temperatures^81^. Yet when adult lifespans are measured at 20°C, larval rearing temperature can have significant impacts on longevity, with the larvae reared in warmer temperatures (25°C), having longer lived adulthoods when moved to 20C post molt^82^. As such, adoption of microfluidic devices for *C. elegans* research will benefit from a concomitant adoption of rearing protocols that avoid confounding environmental changes.

### Fluid flow, feeding, and chip placement

The bacterial solution (NGM supplemented with PXKR1) was pressurized, and flow was maintained at ∼0.5 psi. Six polyethylene tubes 1.5 mm internal diameter (Scientific Commodities Inc) of ∼1 foot led from the bottle to a set of 5 µm syringe filters. The syringe filters and tubing were attached to Y-connector luer stopcocks. Flow could be turned on and off if a particular filter needed changing. The Y shaped system also allowed the researchers to prevent air bubbles by priming the tubing before inserting it into the microfluidic chip. A second set of tubing ∼1 foot continued downstream from the filters and led into the perfusion port of the chip. Each set of 16 arenas shared an entrance and exit port, thus 2 entrance/exit tubes for each chip. A third set of tubing ∼ 1 foot led from the Egg-Counter exit into a resistor chip. Adding resistance into the perfusion system allowed for finer control of embryo speed optimizing capture and detection for later processing. Finally, a 4th set of tubing led out of the resistor chip and into a waste container. The first set of tubing, just above the syringe filter was bundled together and agitated with a small vibration motor. The growth arenas were aligned above the TEC and the temperature evenly distributed to all worms by thermal paste. The imaging arena was centered over the camera and held in place with gaffers’ tape and then secured firmly with hot glue.

### Statistical Analyses

Effects of temperature treatments were tested using a non-parametric Wilcoxon test. For cross-classified frequency data a continuity corrected Yates test was used.

## Conclusions

The development of a microfluidic Egg-Counter for studying *Caenorhabditis* nematode egg-laying behavior exhibits great potential for generating and testing hypotheses regarding the effects of environmental perturbations on a powerful behavioral model.

## Author Contributions

Stephen A. Banse – project administration, formal analysis, visualization, software, methodology, writing, review & editing. Cody M. Jarrett – software, methodology, visualization, writing, review and editing. Kristin J. Robinson – investigation, methodology, writing, review and editing. Benjamin W. Blue – methodology, review and editing. Emily L. Shaw – investigation, review & editing. Patrick C. Phillips – funding acquisition, project administration, methodology, supervision, review and editing.

## Conflicts of interest

The authors have no conflicts to declare.

## Acknowledgements

*C. elegans* strains were provided by the CGC, which is funded by NIH Office of Research Infrastructure Programs (P40 OD010440). We would like to thank Shripad Tuljapurkar, Shawn R. Lockery, and the Phillips lab members for helpful discussion and the University of Oregon’s Lokey Laboratory nanofabrication facility for equipment and technical assistance. Photography services were kindly provided by Derek Robinson. This work was funded by research grants from the National Institute on Aging (R21 AG043988 and R01 AG049396) and the National Institute of General Medical Sciences (R35 GM131838) awarded to PCP.

## Notes

https://doi.org/10.6084/m9.figshare.c.6122667

https://doi.org/10.6084/m9.figshare.c.6122682

https://doi.org/10.6084/m9.figshare.c.6122634

### Competing Interest Statement

The authors have declared no competing interest.

