## Supplemental Figures for "The Egg-Counter: A novel microfluidic platform for characterization of *Caenorhabditis elegans* egg-laying"

### Supplemental Materials

Extensive supplemental materials are available online in three associated collections [1–3] to assist those readers wishing to implement the Egg-Counter platform. In addition to those collections, we provide the following supplemental figures for the general readership.


| | | Page |
|---|---|---|
| Supplemental figure 1 | Egg-Counter software for experimental runs (Egg-Vid-Get) | 2 |
| Supplemental figure 2 | Egg-Counter movement detection | 3 |
| Supplemental figure 3 | Egg-Counter computer aided egg annotation (Egg-Vid-Anno) | 4 |
| Supplemental figure 4 | Cutoff for within versus between bout intervals | 5 |
| Supplemental figure 5 | Egg-Counter CAD drawing | 6 |
| Supplemental figure 6 | Resistance Chip CAD drawing | 7 |
| Supplemental Material References | | 8 |

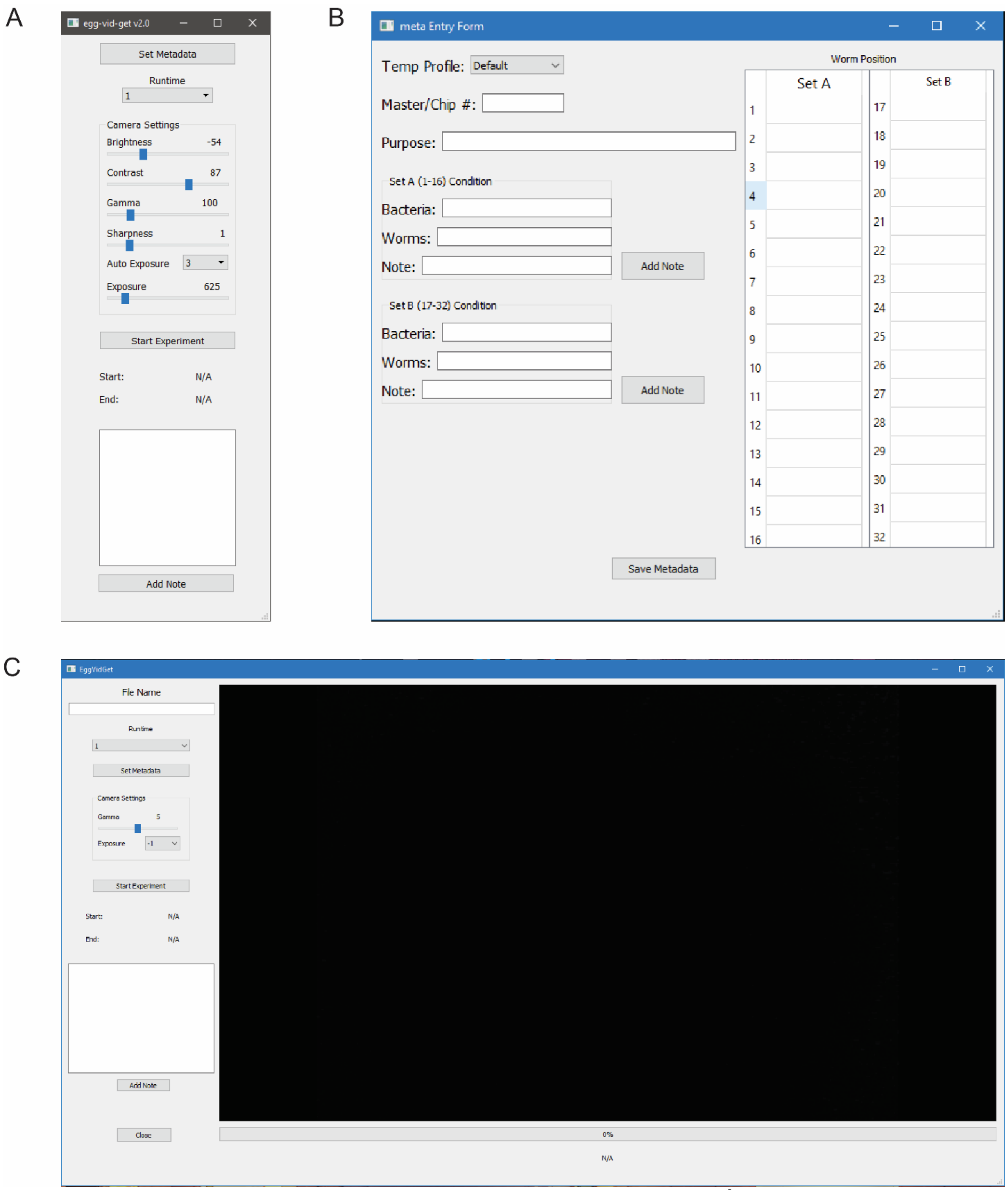


**Supplemental figure 1. Egg-Counter Temperature**. The Egg-Vid-Get software (available in the online collection[2]) provides the necessary functions for running an experiment on the Egg-Counter. Upon launching the software, (A) The provided GUI enables the experimenter to set the image acquisition parameters, enter experimental metadata, and initiate the experiment. When selecting the “Set Metadata” option, a (B) GUI is launched that allows the user to enter experimental metadata for the egg laying experiment. When selecting the “Start Experiment” button, a (C) window is provided with a real-time presentation of the experiment in progress.

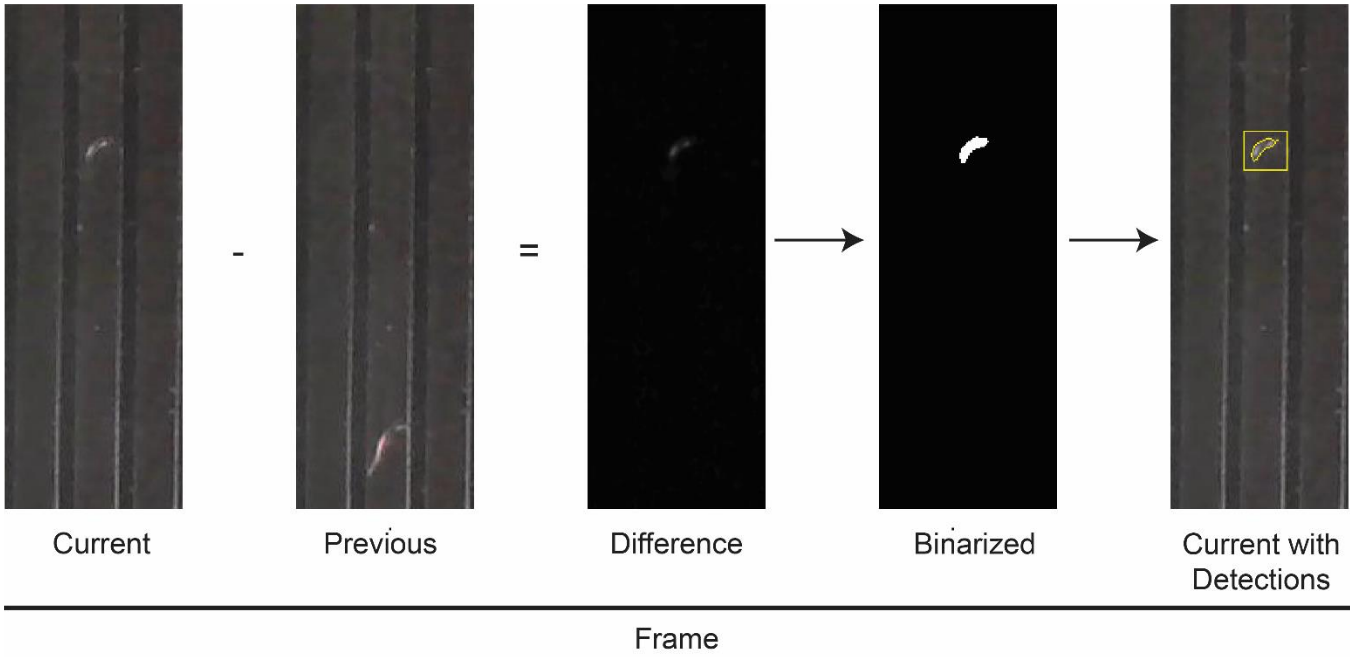


**Supplemental figure 2** The Egg-lane-proc software (available in the online collection[2]) and the egg-vid-get software determine which video frames contain objects of interest. In general, the process uses sequential frames to identify areas of change by simple difference determination. For assembling the list of egg-like objects for use in the Egg-Anno software, several parameters to minimize false positives and streamline the annotation (see online collection[2] for software details).

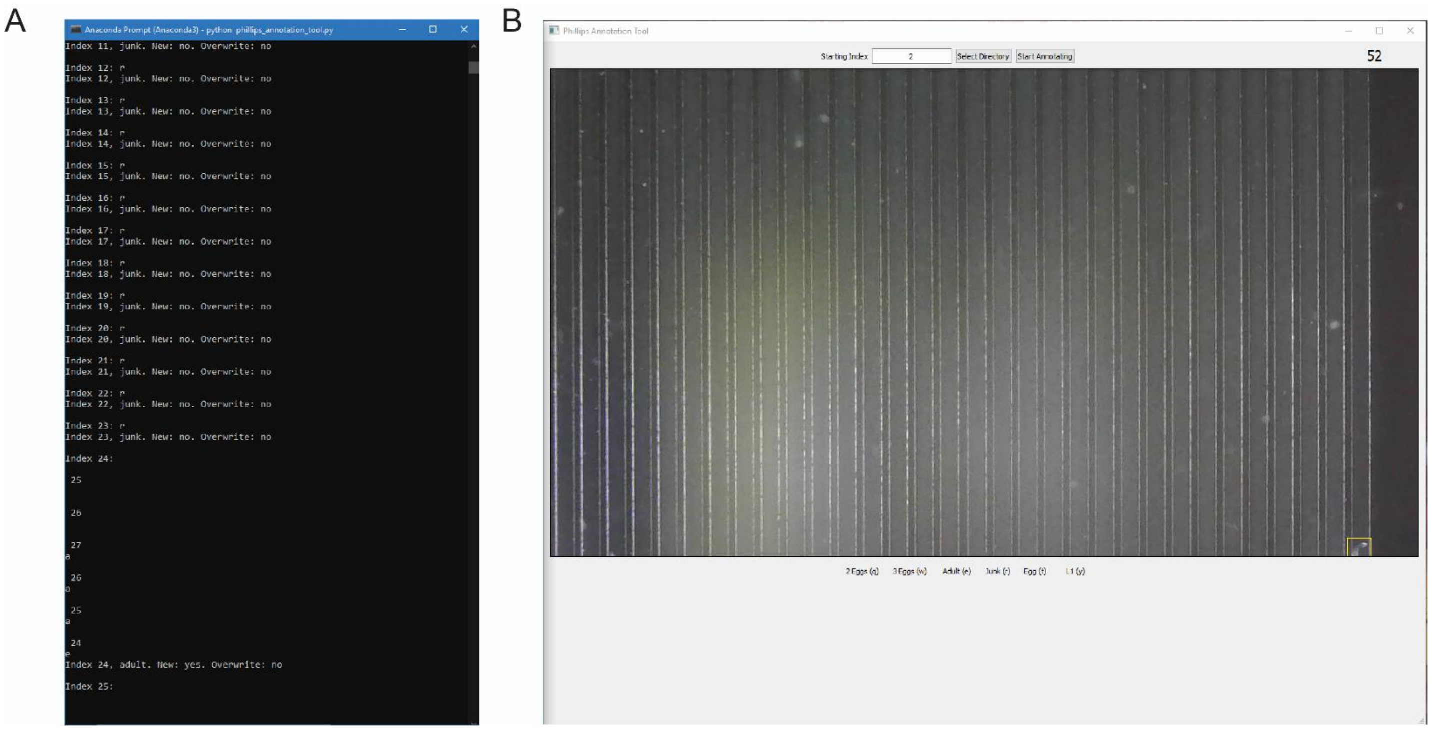


**Supplemental figure 3. The Egg-Counter computer aided annotation.** The Egg-Vid-Anno software (available in the online collection[2]) provides the necessary functions for processing the movie files after an experiment on the Egg-Counter is completed. Upon launching the software, (A) a command window enables the annotation of each egg-like object that was detected by the Egg-Ana software. To make determinations, the user is presented with each frame with the detected objects noted (B). With each entry, the annotations are saved to a post-analysis dataset file.

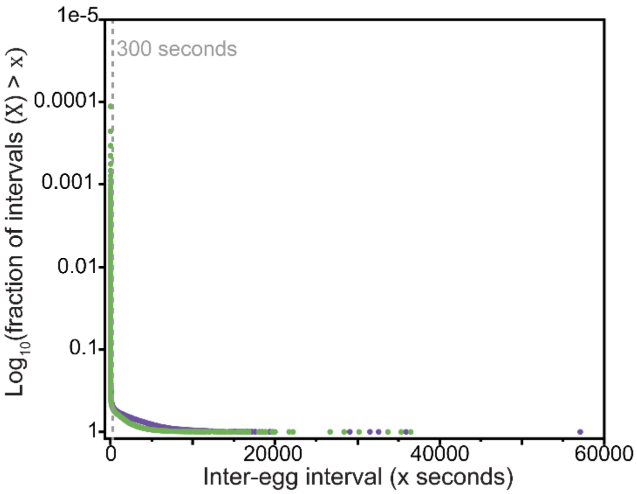


**Supplemental figure 4. Cutoff for within versus between bout intervals.** The log-tail interval distribution plots for the inter-egg-intervals at 15°C (purple) and 20°C (green) provide a visualizable transition between the two classes of intervals, as previously published[4,5]. A transition is observable at ~300 seconds, well between the previously published mean inter-bout-interval of ~20 minutes and the published mean within-bout-interval of ~20 seconds[4,5]. We therefore selected 300 seconds as a cutoff for separating inter-bout and within-bout intervals.

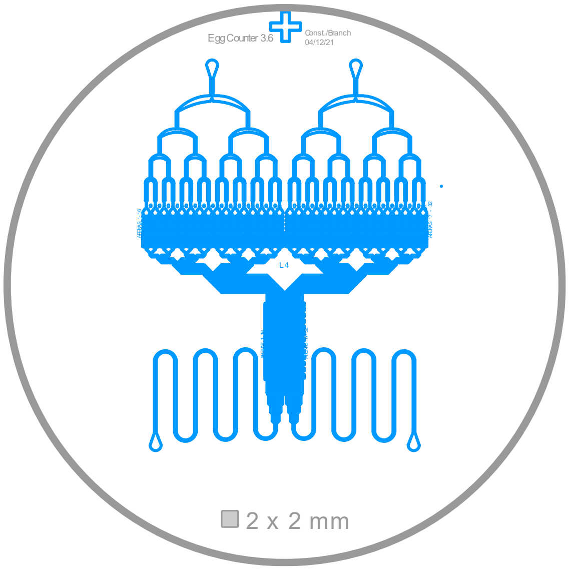


**Supplemental figure 5. A novel microfluidic Egg-Counter**.

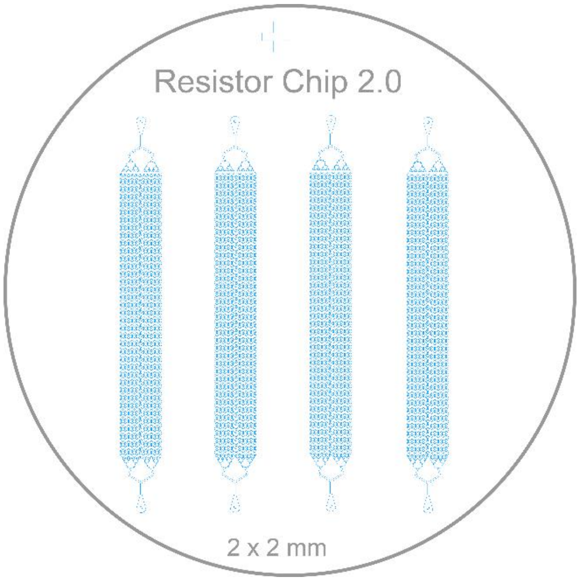


**Supplemental figure 6. Egg-Counter Resistance Chip**
